# CaMKII Phosphorylation of RYR2 is Essential for Arrhythmia in CPVT

**DOI:** 10.1101/2025.09.15.676430

**Authors:** Sofía de la Serna Buzón, Xiaoting Wang, Kristina Chambers, William T. Pu, Vassilios J. Bezzerides

## Abstract

**Background:** Gain of function variants (GOF) in the intracellular calcium (Ca^2+^) release channel *RYR2* predominately underlie the inherited arrhythmogenic syndrome catecholaminergic polymorphic ventricular tachycardia (CPVT). Patients with CPVT are susceptible to life-threatening ventricular arrhythmias triggered by emotional or physical stress. Adrenergic stimulation activates calcium/calmodulin-dependent protein kinase II (CaMKII), which phosphorylates RYR2 on serine 2814 to enhance Ca^2+^ release. Here we assessed the in vivo requirement of this phosphorylation event to unmask the latent arrhythmic potential of CPVT-causing variants.

**Methods:** Using multiplex murine genomic engineering, we mutated the CaMKII phosphorylation site RYR2-S2814 site to alanine (S2814A) in conjunction with a novel CPVT GOF variant (RYR2-R4650I). We systematically interrogated the consequences of CaMKII phosphorylation site ablation on the same (cis) or opposite (trans) allele with the CPVT variant at multiple phenotypic levels.

**Results:** The novel GOF variant *Ryr2-R4650I* conferred a risk of adrenergically inducible ventricular arrhythmia, including bi-directional ventricular tachycardia (BiVT). Ablation of the CaMKII RYR2-S2814 phosphorylation site on the same allele as the GOF variant completely abrogated inducible arrhythmia. In contrast, ablation of this site on the opposite allele did not significantly alter in frequency, type, or length of inducible ventricular tachycardia compared to animals heterozygous for the *Ryr2-R4650I* variant alone. Furthermore, allelic concordance of *Ryr2-R4650I* and *Ryr2-S2814A* did not inhibit adrenergically-induced enhancement of RYR2-dependent functions, including heart rate response, augmentation of cellular Ca^2+^ flux, or intracellular Ca^2+^ release.

**Conclusions:** Our data strongly support the hypothesis that CaMKII phosphorylation of RYR2 at S2814 is necessary and sufficient to unmask the arrhythmogenic phenotype in CPVT. Furthermore, pro-arrhythmia caused by CaMKII-S2814 phosphorylation in CPVT is an *intra-molecular* event, with implications for therapeutic interventions.

## Introduction

The precise regulation of calcium (Ca^2+^) flux within the cardiomyocyte is critical for normal excitation-contraction coupling, with disruption of Ca^2+^ handling underlying many cardiovascular disorders, including arrhythmias and heart failure^1^. During each cardiac cycle, cardiomyocyte depolarization triggers Ca^2+^ influx through T-tubule localized L-type Ca^2+^ channels (LTCCs), dramatically increasing the cytoplasmic Ca^2+^ levels juxtaposed to ryanodine receptor 2 (RYR2) channels, localized on the sarcoplasmic reticulum. This rise in Ca^2+^ triggers RYR2 channel opening and Ca^2+^ release from the sarcoplasmic reticulum, a process known as calcium-induced calcium release. As the major determinant of Ca^2+^ dynamics in cardiomyocytes, the structure and function of RYR2 are central to normal cardiac physiology and are affected in both genetic and acquired disease states^2^.

The RYR2 channel is a massive 2.2 megadaltons homo-tetramer, with each human monomer comprising 4967 amino acid residues. This channel exhibits radial symmetry, with each monomer contributing equally to the central pore. High-resolution structural analysis over the last two decades has indicated that RYR2 channels resemble a “mushroom” with a large shell-like cytoplasmic facing region and a more narrow channel or “stalk” crossing the SR membrane^3^. The direct binding of associated proteins and a diverse array of post-translational modifications across the cytoplasmic shell regulate channel gating and Ca^2+^ release. Therefore, RYR2 dysfunction may be a result of dysregulated extrinsic signaling pathways, gain or loss of function (GOF, LOF) variants, or a combination of both factors^4^. Despite intensive research, the relative contribution of extrinsic and intrinsic factors in RYR2 regulation is not fully understood.

RYR2 channel dysfunction is implicated in both genetic and acquired forms of heart disease, highlighting its critical role in maintaining cardiac Ca^2+^ homeostasis^5^. Indeed, single missense gain-of-function (GOF) variants in RYR2 are responsible for the inherited arrhythmia syndrome catecholaminergic polymorphic ventricular tachycardia (CPVT)^6^. At baseline, patients with CPVT do not display structural abnormalities or baseline ECG changes but some patients present with sinus bradycardia^7^. Only in the presence of β-adrenergic stimulation is the latent effect of the RYR2 pathogenic variant unmasked, leading to increased diastolic Ca^2+^, delayed afterdepolarizations (DADs), and electrical instability^8^. β-adrenergic stimulation of cardiomyocytes activates Ca^2+^/calmodulin-dependent protein kinase II (CaMKII), which enhances Ca^2+^ handling by phosphorylating multiple targets, including RYR2 at serine 2814 (S2814). RYR2 phosphorylation on this residue increases the channel’s open probability^9,10^. We previously reported that homozygous ablation of this phosphorylation site in iPSC-CMs harboring a CPVT-causing variant (*RYR2^R4651I/+^*) prevented CPVT-associated arrhythmias^8^. However, these studies were limited to iPSC-CM models and did not investigate the requirement of CaMKII phosphorylation in the same allele or the opposite allele to the CPVT variant.

To overcome these limitations and address the structural context of RYR2 regulation, we generated allele-specific murine models by multiplexed genome editing. By pairing the CPVT-causing R4650I variant (analogous to human R4651I) with the S2814A phosphorylation-null mutation in *Cis* (same allele) and *Trans* (opposite allele) configurations, we interrogated the intra-versus inter-monomer mechanisms governing RYR2-mediated arrhythmogenesis across functional scales, including whole-animal electrophysiology, cardiomyocyte calcium imaging, and channel-level phosphorylation analysis.

## Methods

### Generation of mouse lines

All experimental animals were housed in the Boston Children’s Hospital animal facility under institutionally approved IACUC protocols, and littermates of both sexes were used for all experiments. For phenotypic outcomes or single cell outcomes the researcher performing the experiment and/or analysis was either blinded to the genotype or another researcher randomized the phenotypic outcomes prior to analysis with unblinding only after the analysis was complete.

Mouse models were generated using CRISPR/Cas9-mediated genome editing in C57BL/6 embryos. Guide RNAs (gRNAs) and homology-directed repair (HDR) templates (Table 1) were designed to introduce single point mutations at the RYR2 locus. For the RYR2-S2814A mutation, serine 2814 was substituted with alanine (S2814A) to prevent CaMKII phosphorylation. A silent mutation creating a restriction enzyme site was also simultaneously added to facilitate genotyping. The homozygous RYR2-S2814A line, referred to as RYR2-Unphos, was independently generated to serve as a control for complete phosphorylation loss. The RYR2-R4650I mutation was introduced by replacing arginine 4650 with isoleucine, corresponding to the human RYR2-R4651I CPVT variant^8,11^. A mixture of active gRNA transcripts, single-stranded oligodeoxynucleotide donor, and Cas9 mRNA was microinjected into the cytoplasm of C57BL/6 embryos, which were then implanted into pseudo pregnant females. Founder mice were screened for knock-in mutations by Sanger sequencing for validation. Only heterozygous *Ryr2^R4650I/+^* (*Ryr2-CPVT*) mutants were studied, as generally only heterozygous GOF variants are observed in CPVT patients. To obtain the RYR2-cis-coupled double mutant *Ryr2^R4650I;S2814A/+^* embryos were simultaneously injected with gRNAs for both mutations, and offspring were screened for the presence of both mutations. Backcrosses to wild type mice confirmed that the mutations reside in the same allele. The RYR2-trans-double mutant *Ryr2^R4650I/S2814A^* (RYR2-*Trans)* was generated by crossing the *Ryr2-CPVT* and *Ryr2-Unphos* lines, ensuring segregation of mutations onto opposite alleles. All mouse lines were backcrossed to C57BL/6J for six generations to establish stable lines.

### Induction of arrhythmia in transgenic mouse models

The assessment of arrhythmia inducibility by drug challenge was performed on animals at 2 to 4 months of age. Mice were anesthetized with isoflurane (3% induction, 1.5% maintenance in oxygen) and then transferred to a heated platform and maintained via a nose cone. Adequate sedation was confirmed by the absence of reflex response to toe pinching before procedure initiation. A two-lead surface electrocardiogram (ECG) (leads I and II) was recorded using subcutaneous needle electrodes placed in the right forelimb, left forelimb, and left hindlimb. Baseline ECG parameters were recorded for at least 3 minutes before pharmacological challenge.

To assess arrhythmia susceptibility, mice underwent sequential administration of adrenergic agonists and caffeine. A single intraperitoneal (IP) injection of isoproterenol (2 mg/kg) was administered to enhance β-adrenergic stimulation. ECGs were continuously recorded for 3 minutes post-injection. After isoproterenol, mice received a combined IP injection of epinephrine (4 mg/kg) and caffeine (120 mg/kg) to potentiate Ca^2+^-mediated arrhythmogenesis. ECGs were recorded for an additional 3.5 minutes post-injection. Heart rates were recorded after each drug stimulation and compared to baseline heart rate levels. For analysis, PVCs, single couplets, BiVT and VT during this recording period were considered a positive response. Ectopy rate is calculated as the proportion of abnormal beats to total beats captured after the characteristic epinephrine/caffeine heart rate dip. The longest duration of ectopic beats was defined as the longest period of abnormal beats during the three and a half minutes captured post stimulation.

### Histology

Tissue samples were fixed in 4% PFA in PBS, embedded in paraffin, and sectioned at 5 µm. Sections were deparaffinized, rehydrated, and stained using Masson’s Trichrome according to the manufacturer’s protocol to visualize collagen deposition. Briefly, sections were incubated in Bouin’s solution at 56°C for 1 hour, followed by staining with Weigert’s iron hematoxylin, Biebrich scarlet-acid fuchsin, phosphotungstic/phosphomolybdic acid, and aniline blue. Collagen fibers stained blue, muscle fibers stained red, and nuclei appeared black. Images were captured using a brightfield microscope, and sections were evaluated for fibrotic presentation using ImageJ software.

### Echocardiography

Non-invasive assessment of ventricular performance was performed using the VEVO F2 (Visual Sonics) on awake mice. After ventral hair removal, a 30 MHz echo probe was used to obtain parasternal long and short axes to ensure on-axis imaging. While in the parasternal short view, 15 second M-mode imaging clips through the left ventricle (LV) were obtained to directly measure functional parameters in both systole (s) and diastole (d): LV interventricular septal thickness (IVS), LV internal dimension (LVID), and posterior wall thickness (PW)^12^. The M-mode images were generated at the level of the papillary muscles. Fractional shortening (FS) was calculated using the following formula:

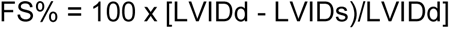

Any animal with a heart rate less than 400 bpm was either excluded from analysis or was analyzed on a subsequent day. At least four measurements were acquired and averaged for each animal at each time point.

### Adult cardiomyocyte isolation and calcium imaging

Knock-in mice and their wild-type littermates of either sex and 8–12 weeks-old were used for the isolation of ventricular cardiomyocytes. Ventricular cardiomyocytes were isolated by retrograde perfusion through the aorta using a standardized enzymatic digestion protocol with minor modifications^13^. Briefly, mice were injected with heparin before euthanasia. Hearts were extracted, perfused and digested with collagenase II (Worthington) using Langendorff apparatus at 37°C. Once the cardiomyocytes were properly dissociated, the hearts were removed from the cannula, gently agitated with a pair of tweezers, and strained using a 100 µm cell strainer. Cells were resuspended in 50 μM CaCl_2_ Tyrodes with 10 mM 2, 3-Butanedione monoxime (BDM) and 10 mM taurine and allowed to settle to the bottom by gravity for 15 minutes. Intracellular Ca^2+^ storage was gradually restored by replacing the buffer with Tyrodes containing increasing concentrations of CaCl_2_ (400 μM, 900 μM, and 1.2 mM, respectively) every 4 minutes. Once recalcification was complete, cardiomyocytes (CMs) were plated to laminin-coated coverslips.

### Whole cardiomyocyte calcium imaging

For whole-cell calcium imaging and contractility measurements, adult CMs were incubated with 5μM Fura-2 (Invitrogen, Cat# F1221) in Tyrode’s buffer for 20 min prior to imaging. Individual cells were identified as rod-shaped CMs and a mask in the optical path was centered around each cell to measure Fura-2 fluorescence at 515 nm with alternating 340 and 380 nm excitation. Cells were electrically paced at 1Hz (10-15 mA, 8 msec square wave pulses) for 30 seconds followed by cessation of pacing and continued recording for an additional 60 s. Recordings were repeated after the addition of isoproterenol (100 nM). Post pacing events (PPEs) were recorded after the cessation of pacing and expressed as the number of spontaneous triggered Ca^2+^ transients and waves during the recording period. Whole-cell calcium imaging experiments were performed with the investigator blinded to the genotype of the mice to reduce bias in data acquisition and interpretation. Contraction and Ca^2+^ transient parameters were calculated using IonWizard (IonOptix).

### Measurement of calcium sparks

To measure spontaneous Ca^2+^ sparks in adult cardiomyocytes, isolated cells underwent Ca^2+^ re-introduction and then stained with Fluo4-AM (Invitrogen, Cat# F14201) at a concentration of 20 μM at 37 C for 5 minutes. Cardiomyocytes were washed once with Tyrode’s solution containing in mM: 140 NaCl, 4 KCl, 10 HEPES, 2 Sodium pyruvate, 10 glucose, at pH 7.4. Cells were plated on laminin-coated glass coverslips and imaged by confocal line scanning on an LSM880 (Zeiss corporation) with a 40x oil immersion objective. Repeat line scan time was 4.1 μs with a spatial resolution of 0.083 μm, excitation wavelength of 488 nm and a pinhole diameter of 2 μm. The frequency of Ca^2+^ sparks and spark mass were calculated by a program written in IDL^14^.

### Statistics

Statistical analyses were performed with GraphPad Prism. Unless otherwise stated, values are presented as the mean plus and minus the standard error of the mean (± SE). In general, we applied an ordinary one-way ANOVA with Tukey’s post hoc test to model the data; however, in instances where variance was unequal across groups, we used Welch’s ANOVA with Dunnett’s T3 post hoc test. To compare ectopy rates across the five experimental groups, we employed a nonparametric Kruskal– Wallis test, as the data were right-skewed due to a large number of animals exhibiting no ectopy (i.e., zero events). For analyses of Ca^2+^ spark frequency and post-pacing events, which consisted of over dispersed count data with many zeros, we used generalized linear models with a negative binomial distribution and log link, followed by post hoc pairwise contrasts estimated with the emmeans package in R (R version 4.1.3). or Ca^2+^ transient parameters (peak amplitude, upstroke velocity, downstroke velocity), we used generalized linear mixed-effects models with a Gamma distribution and log link, including random intercepts for cell line and individual cells to account for repeated measures. Model significance was assessed using Wald chi-square tests, and post hoc pairwise contrasts were adjusted for multiple testing (Tukey or Holm as specified).

## Results

### Generation of a novel CPVT mouse model

Over 200 *RYR2* pathogenic variants have been reported to cause CPVT in humans^15^. These mutations cluster in four distinct regions (I-IV) with correspondingly diverse mechanisms of channel dysfunction^16^. We previously identified a novel GOF *RYR2* variant p. (Arg4651Ile) in a CPVT patient with recurrent atrial and ventricular arrhythmias^8^. This variant demonstrated abnormal Ca^2+^ handling in induced pluripotent stem cell (iPSC) models, but a direct link to arrhythmia was missing^8^. To determine the effects of this GOF variant in an animal model, we introduced the murine orthologue (R4650I) into the *Ryr2* gene by CRISPR/Cas9 genome editing (Figure 1A). To confirm the lack of structural heart disease or changes in ventricular performance^17^, we performed echocardiography of *Ryr2^R4650I/WT^*and wild-type littermates. Similar to CPVT patients, left ventricular (LV) fraction shortening LV end diastolic volumes (LVEDVs) were indistinguishable between *Ryr2^R4650I/WT^* and WT animals (Figure 1B, C). Stained histologic sections from *Ryr2^R4650I/WT^* hearts also did not show significant fibrosis consistent with the lack of any structural abnormalities (Figure 1D). Continuous ECG recording under anesthesia showed that baseline ECGs were comparable between genotypes (Fig. 1E-G). However, adrenergic stimulation with isoproterenol (ISO) followed by epinephrine and caffeine (EPI/CAFF) induced frequent premature ventricular contractions (PVCs) and runs of ventricular tachycardia (VT), including bidirectional tachycardia (BiVT) (Figure 1E-G). Ventricular tachycardia was noted in 7/9 (77.8%) of *Ryr2^R4650I/WT^*animals (Figure 1G). These data demonstrate that RYR2-R4650I, the murine ortholog of human RYR2-R4651I, is a new CPVT model characterized by adrenergically-inducible ventricular arrhythmia in the absence of structural disease.

**Figure 1.**
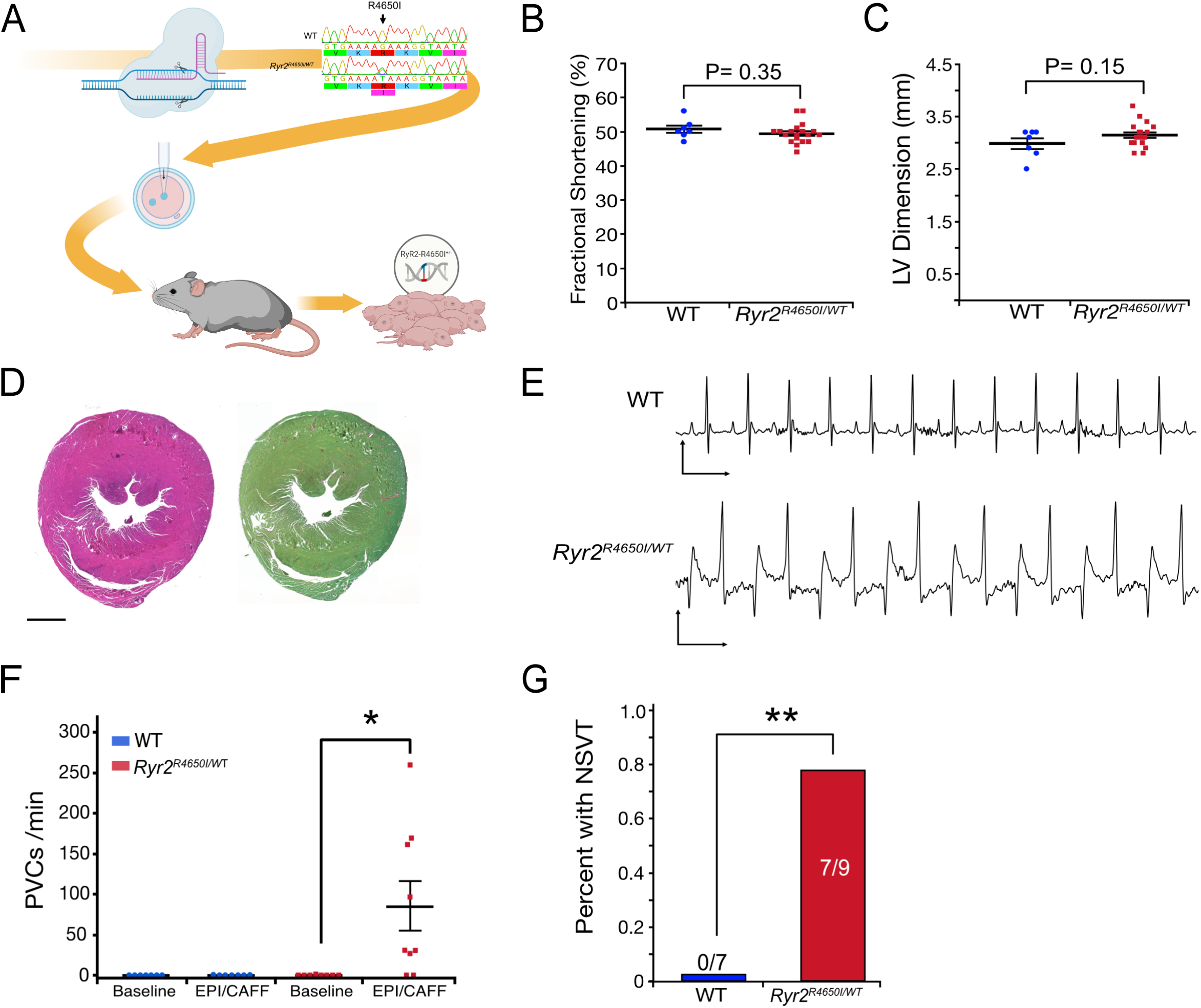
Generation and characterization of a novel CPVT mouse model with the Ryr2^R4650I/WT^ mutation. **A,** Injection of a specific guide RNA, donor DNA template, and purified Cas9 protein into mouse zygotes was used to generate a knock-in mutation of arginine to isoleucine at position 4650 (R4650I) in the Ryr2 gene. This mutation corresponds to the human RYR2-R4651I variant associated with catecholaminergic polymorphic ventricular tachycardia (CPVT). Founder pups were screened via Sanger sequencing to confirm successful editing. **B,** Fractional shortening (FS), measured by non-invasive echocardiography, showed no significant differences between wild-type (WT) and Ryr2^R4650I/WT^ mice, consistent with preserved ventricular function (Student’s *t* test). **C,** Left ventricular internal diameters in systole also did not differ significantly between genotypes, indicating no change in chamber size (Student’s *t* test). **D,** Histologic sections of Ryr2^R4650I/WT^ hearts stained with hematoxylin and eosin (H&E, left) and Sirius red (right) revealed no evidence of fibrosis or cardiomyopathy. Scale bar = 1 mm. **E,** ECG recordings performed after intraperitoneal injection of epinephrine (4 mg/kg) and caffeine (120 mg/kg) demonstrated normal sinus rhythm in WT mice (top) and bidirectional ventricular tachycardia (BiVT) in Ryr2^R4650I/WT^ mice (bottom), a hallmark of CPVT. **F,** Quantification of premature ventricular contractions (PVCs) per minute was performed during three-minute epochs at baseline and after epinephrine/caffeine (EPI/CAFF) stimulation. Only Ryr2^R4650I/WT^ mice exhibited a statistically significant increase in PVCs following stimulation (Steel-Dwass test, *P* < 0.05). **G,** Incidence of non-sustained ventricular tachycardia (NSVT), defined as runs of four or more consecutive beats, was assessed following an adrenergic challenge. NSVT occurred in 0 of 7 WT mice and in 7 of 9 Ryr2^R4650I/WT^ mice (Fisher’s exact test, P < 0.01). Significance values: * P<0.05, ** P<0.01, *** P <0.005, **** P <0.001.

### Ablation of the CaMKII phosphorylation site in combination with a CPVT variant

Despite normal baseline ECGs, CPVT patients carry a latent arrhythmogenic risk that is only unmasked under adrenergic stimulation in the presence of a pathogenic variant ^6,8^. To directly investigate the molecular hierarchy of CaMKII phosphorylation and RYR2 dysfunction, we generated three additional mouse lines incorporating *Ryr2-R4650I* and/or the ablation of the CaMKII phosphorylation site (*Ryr2-S2814A*) in several configurations, generating five experimental groups: (1) wild type (WT), (2) the CPVT mutant *Ryr2^R4650I/WT^* referred to as *Ryr2-CPVT*, (3) the homozygous CaMKII phosphorylation mutant *Ryr2^S2814A/S2814A^* referred to as *Ryr2-Unphos*, (4) the cis-coupled double mutant with both variants on the same allele, *Ryr2^R4650I;S2814^*^A/WT^ referred as *Ryr2-Cis*, and (5) the trans-double mutant where the variants are on the opposite alleles *Ryr2^R4650I;S2814A^*, referred to as *Ryr2-Trans* (Figure 2A). Echocardiographic analysis of conscious mice across these genotypes demonstrated that none of the edits negatively affected ventricular function (Figure 2C). There were no significant differences in fractional shortening (FS, *P* = 0.298; ANOVA: F = 1.269) intraventricular systolic dimension (IVs, *P* = 0.213; ANOVA: F = 1.523), or left ventricular posterior wall dimension (LVPWd, *P* = 0.462; ANOVA: F = 0.919) indicating preserved systolic function across all genotypes (Figure 2C).

**Figure 2.**
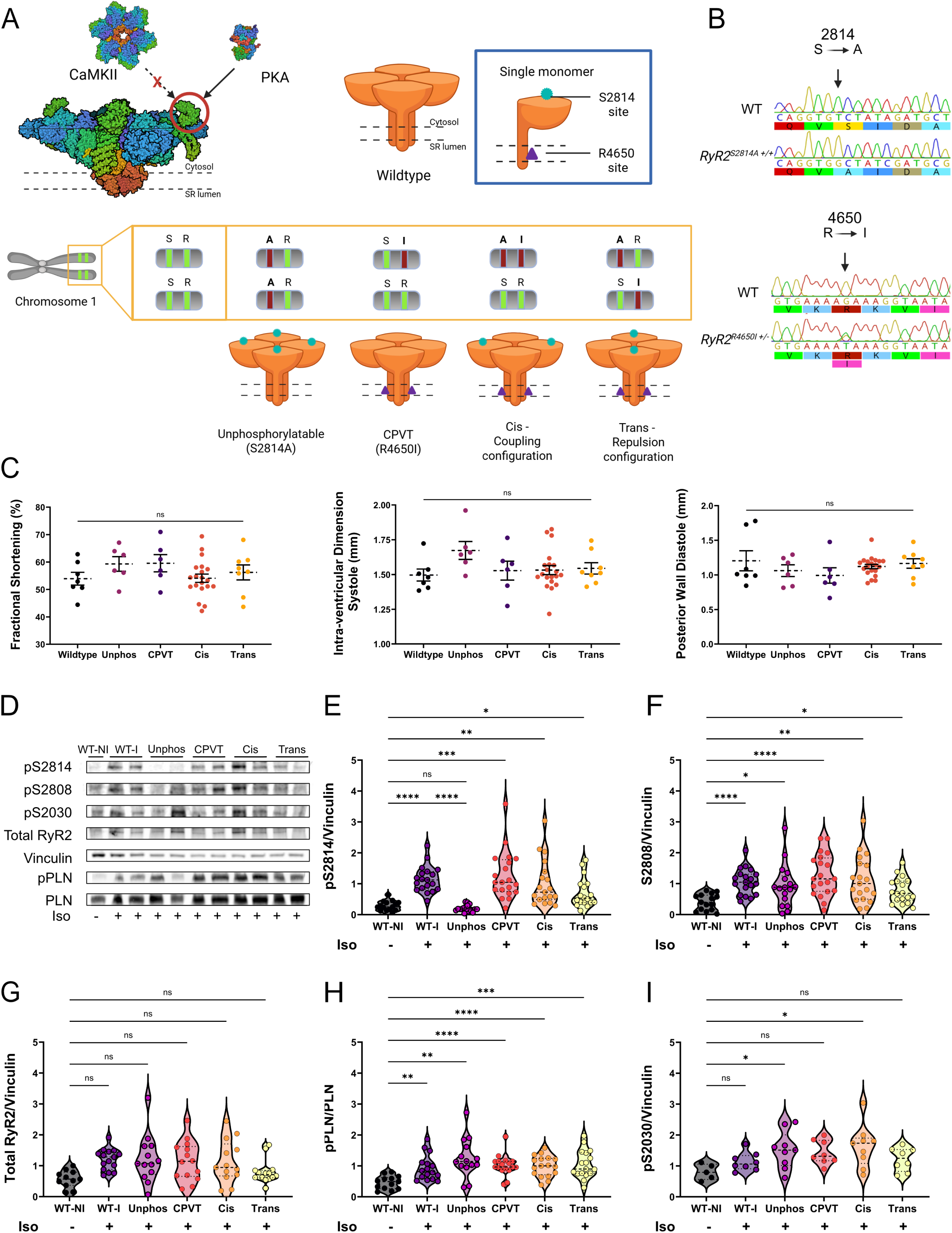
Baseline characterization of RYR2 mouse models with distinct combinations of R4650I and S2814A mutations. **A,** Diagram representing the five mouse genotypes used in this study, based on combinations of two mutations in the RYR2 gene: the CPVT-linked R4650I mutation and the non-phosphorylatable S2814A mutation. Image created with Biorender.com. **B,** Sanger sequencing confirmation of the S2814A and R4650I mutations in genomic DNA from the engineered mouse lines. **C,** Echocardiographic assessment of cardiac structure and function. Graphs display fractional shortening, right; intra-ventricular dimension in systole, middle; and posterior wall thickness in diastole, left across all five genotypes. No statistically significant differences were observed compared to wildtype, indicating preserved baseline cardiac function and structure. Statistical analysis was performed using one-way ANOVA with Dunnett’s multiple comparisons test. Sample sizes ranged from n = 6 to 20 per group. **D,** Representative western blots corresponding to each quantitative analysis, including blots for pS2814, pS2808, pS2030, total RYR2, vinculin, and pPLN/PLN. Western blot analyses of RYR2 phosphorylation and protein levels across six experimental groups: wildtype without isoproterenol, and wildtype, *Unphos*, *CPVT*, *Cis*, and *Trans* genotypes, all treated with 4mg/kg isoproterenol prior to sacrifice. Protein was isolated from whole heart lysates. **E,** Phosphorylation at RYR2-S2814 (pS2814), normalized to vinculin, was significantly increased in all isoproterenol-stimulated groups compared to wild type without isoproterenol (*P* < 0.05). Statistical analysis: Welch’s ANOVA with Dunnett’s T3 multiple comparisons test (W = 28.54, DFn = 5, DFd = 45.15, *P* < 0.0001) ; n = 15–20 per group. **F,** Phosphorylation at RYR2-S2808 was significantly increased in all stimulated groups compared to wild type without isoproterenol (*P* < 0.05), except in the *Trans* group. Statistical analysis: Welch’s ANOVA and Dunnett’s T3 post hoc analysis: W = 10.66, DFn = 5, DFd = 47.11, *P* < 0.0001; n = 15–20 per group. **G,** Total RYR2 levels normalized to vinculin were unchanged across all groups, indicating that the mutations did not affect overall RYR2 expression. Statistical analysis: one-way ANOVA with Tukey’s test (*F* = 2.910, *P* = 0.0188); n = 11–15 per group. **H,** Phospholamban phosphorylation (pPLN) normalized to total phospholamban was significantly increased in all stimulated groups compared to wild type without isoproterenol. Statistical analysis: Welch’s ANOVA and Dunnett’s T3 post hoc analysis (W = 11.05, DFn = 5, DFd = 40.36, *P* < 0.0001) n = 13-17 per group. **I,** Phosphorylation at RYR2-S2030 showed no significant differences except for elevated levels in the *Unphos* and *Cis* groups compared to unstimulated wild type. Statistical analysis: ordinary one-way ANOVA with Tukey’s post hoc test (*F* = 2.590, *P* = 0.0399); n = 5–9 per group. For all tests above, statistical significance was defined as P < 0.05. All analyses were performed using GraphPad Prism version 10.5.0. Significance values: * P<0.05, ** P<0.01, *** P <0.005, **** P <0.001.

β-receptor stimulation activates multiple downstream signaling events, including activation of CaMKII followed by phosphorylation of CaMKII substrates. We performed western blotting on whole heart protein lysates before and after stimulation with isoproterenol (ISO) to quantify ISO-stimulated phosphorylation events. Total RYR2 protein levels were not significantly different across genotypes (Figure 2D,E). To confirm CaMKII activation, we examined adrenergically mediated phosphorylation of phospholamban (PLN) at threonine 17, a known CaMKII target¹⁸. Western blot analysis demonstrated a significant increase in pPLN across all genotypes following isoproterenol stimulation (Figure 2D-F). At RYR2, ISO increased phosphorylation of the CaMKII target S2814 in wild type hearts, and this phosphorylation was abolished by S2814A mutation in RYR2-Unphos mutants (Fig. 2D, G). In contrast, ISO-induced S2814 phosphorylation was preserved in *Ryr2-CPVT*, *Ryr2-Cis* and *Ryr2-Trans* lines (Figure 2D, G). Adrenergic stimulation also induces activation of PKA and hyperphosphorylation of RYR2-S2808, which has also been reported to increase RYR2 channel open probability^18^. We confirmed that ISO increased RYR2-S2808 phosphorylation in all genotypes, including *RYR2-Unphos* (Fig. 2D, H). Collectively, these data validated the *RYR2-Unphos* mutation, demonstrated that ISO-induced phosphorylation at S2814 is preserved in *Ryr2-CPVT*, *Ryr2-Cis* and *Ryr2-Trans* hearts, and showed that perturbation of the S2814 phosphorylation site does not alter PKA-mediated phosphorylation at S2808.

### Characterization of heart rate responsiveness of CPVT mice

More than 50% CPVT patients have bradycardia and sinus node dysfunction even without beta-blocker therapy20. To investigate whether this phenotype is dependent upon RYR2-S2814 phosphorylation in cis- or in trans- to the CPVT-causing variant, we analyzed the heart rate response of our models to isoproterenol. Prior to drug treatment there were no significant differences in resting heart rate among the different genotypes tested (Figure 3B). However, the relative increases in heart rate with ISO stimulation were lower in the *Ryr2-CPVT* and *Ryr2-Trans* animals as compared to WT controls (99.08 vs.148.7, P = 0.0347, 112.3 vs.148.7, P = 0.1319) (Figure 3C). Conversely, heart rate responses to EPI/CAFF were also significantly blunted in the *Ryr2-CPVT* and *Ryr2-Trans* lines (112.5 vs. 237.7, P < 0.001, 133.4 vs. 237.7, P < 0.001; Figure 3D).

**Figure 3.**
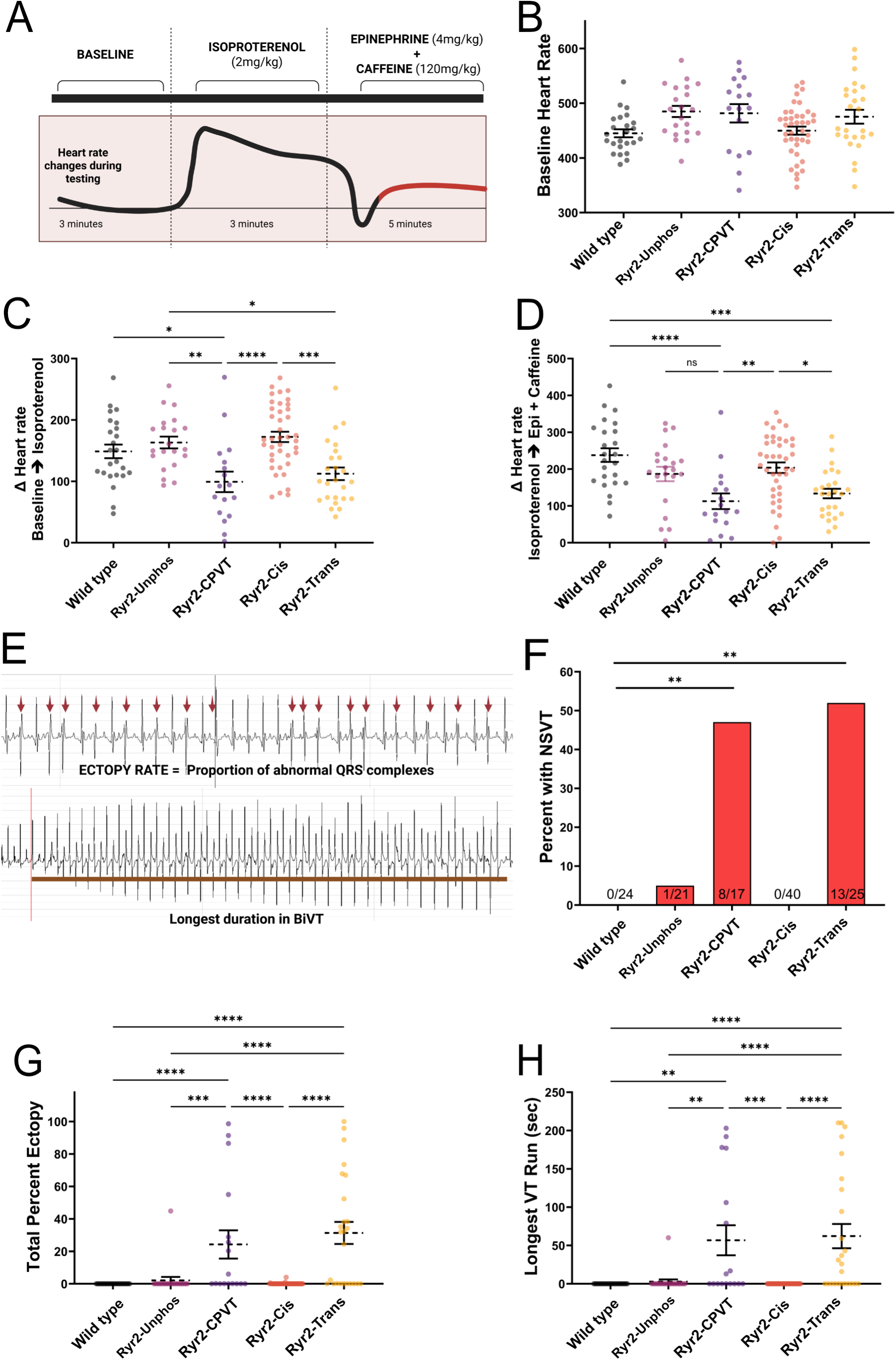
Electrophysiologic Responses and Arrhythmia Burden in Sedated Mice Following Drug Stimulation. **A**, Schematic representation of the sedated in vivo electrophysiology (EP) recording paradigm with drug stimulation protocol. Following isoproterenol (ISO) administration (2mg/kg), there is a stereotypical increase in heart rate followed by a paradoxical drop after administration of epinephrine (4mg/kg) and caffeine (120mg/kg) (EPI/CAFF), with ventricular ectopy typically observed upon the return of heart rate to pre-stimulation levels. **B**, Baseline heart rates did not significantly differ across the five experimental groups. **C**, The change in heart rate was measured from baseline levels following isoproterenol administration. The relative increases in heart rate with ISO stimulation were lower in the *Ryr2-CPVT* and *Ryr2-Trans* animals as compared to either WT or the other lines (99.08 vs. 148.7, *P* = 0.0347, 112.3 vs. 148.7, *P* = 0.1319; ANOVA: F = 8.595, *P* < 0.001; post hoc test: Tukey). **D**, Heart rate response to combined epinephrine and caffeine stimulation relative to isoproterenol maximum. The relative heart rate decrease in response to EPI/CAFF were significantly blunted in the *Ryr2-CPVT* and *Ryr2-Trans* lines (112.5 vs. 237.7, *P* < 0.001, 133.4 vs. 237.7, *P* < 0.001; ANOVA: F = 8.045, *P* < 0.001; post hoc test: Tukey). **E**, Representative ECG tracings illustrating abnormal beats included in the calculation of percent ectopy (top). The second panel (bottom) shows the longest ventricular tachycardia (VT) run, defined as the longest sequence of uninterrupted VT observed during the entire recording session. **F**, Proportion of animals in each group exhibiting non-sustained ventricular tachycardia (NSVT). **G**, Total percent ectopy observed in each group during drug stimulation. To compare ectopy rates across the five experimental groups, we employed a nonparametric Kruskal– Wallis test, as the data were right-skewed since many animals exhibit no ectopy (i.e., zero events) (H = 55.11, df = 4, *P* < 0.0001, post hoc test: Dunn’s). Statistical significance was defined as *α* = 0.05.Sample sizes ranged from *n* = 17 to 40 animals per group. **H**, Longest ventricular tachycardia (VT) run recorded per group. Data are shown as mean ± SEM. Kruskal-Wallis W = 49.72, *P* <0.001; Dunn’s Post hoc test; n=17-40 per group. All analyses were performed using GraphPad Prism version 10.5.0. Significance values: * P<0.05, ** P<0.01, *** P <0.005, **** P <0.001.

### Characterization of adrenergically-induced arrhythmia

Targeted CaMKII inhibition or genetic ablation of its primary phosphorylation site on RYR2 (S2814) reverses the CPVT phenotype in single cells and tissue models^8^. While supportive of the importance of CaMKII phosphorylation of RYR2 at S2814 in the pathogenesis of CPVT, these data were based on the homozygous ablation of S2814 and only in cellular models. To test the importance of RYR2-S2814 phosphorylation for CPVT arrhythmogenesis in intact animals, we acquired continuous ECG recordings during serial administration of ISO followed by EPI/CAFF^19^. ISO induced tachycardia. Subsequent EPI/CAFF treatment transiently depressed HR, which then recovered to a greater than baseline rate. When present, ventricular ectopy and BiVT occurred immediately following this HR recovery (Figure 3A, red line, Figure 3E). Under this adrenergic stimulation protocol, nearly 50% of both *Ryr2-CPVT* (8 of 17) and *Ryr2-Trans* (13 of 25) animals developed non-sustained ventricular tachycardia (NSVT), defined as greater than 3 ventricular beats in a row (Figure 3F). In contrast, NSVT episodes were absent or nearly absent in *WT* (0 of 24), *Ryr2-Unphos* (1 of 21), and *Ryr2-Cis* (0 of 40) mice (Figure 3F).

In addition to episodes of ventricular tachycardia, CPVT patients have increased ventricular ectopy overall^20^ and therefore after stimulation with EPI/CAFF we quantified the percentage of total beats that were ectopic. Again, the frequency of ventricular arrhythmia was significantly higher in *Ryr2-CPVT* and *Ryr2-Trans* animals compared to the other groups (mean ± s.e.: *Ryr2-CPVT* 24.25 ± 8.682; *Ryr2-Trans* 31.28 ± 6.804 versus WT 0.002 ± 0.002; *Ryr2-Unphos* 2.134 ± 2.134 and *Ryr2-Cis* 0.143 ± 0.104; Figure 3G). In addition to increasing the frequency of single PVCs and NSVT, adrenergic stimulation induced continuous, prolonged episodes of VT, including BiVT, in *Ryr2-CPVT* (56.76 ± 19.58 seconds) and *Ryr2-Trans* animals (62.13 ± 15.86 seconds), with only a single prolonged episode of VT in *Ryr2-Unphos* cohort (2.857 ± 2.86 seconds) and none in either WT or *Ryr2-Cis* mice (Figure 3H).

Taken together, these data demonstrate that the RYR2-R4650I CPVT variant encodes a latent propensity to ventricular arrhythmias, which requires RYR2-S2814 phosphorylation on the Cis allele to be unmasked.

### Effect of cis- and trans-RYR2-S2814 phosphorylation on individual CPVT cardiomyocyte physiology

Excessive diastolic Ca^2+^ leak through RYR2 channels is a proposed common mechanism for most forms of CPVT^21^. To determine if the RYR2-R4650I variant causes abnormal Ca^2+^ signaling at the cellular level, we acutely isolated adult mouse cardiomyocytes and loaded them with the ratiometric fluorescent Ca^2+^ indicator Fura-2. We then paced isolated cardiomyocytes at 1 Hz for 1 min, followed by the cessation of pacing for an additional 30 s (Figure 4A). In response to this paradigm, WT cardiomyocytes rarely displayed Ca^2+^ signaling events after the cessation of pacing (Figure 4B and Figure 4C). In contrast, cardiomyocytes isolated from *Ryr2-CPVT* animals displayed abnormal Ca^2+^ post-pacing events (PPEs) (*P* < 0.032) (Figure 4C). The frequency of these events was further augmented by treatment with 100 nM ISO (*P* < 0.026) (Figure 4C). In contrast, homozygous ablation of the CaMKII phosphorylation site (*Ryr2-Unphos*) or incorporation of S2814A along with R4650I (*Ryr2-Cis*), completely abrogated post-pacing PPEs even under adrenergic stimulation (Figure 4B and Figure 4C). Cardiomyocytes from *Ryr2-Trans* animals displayed increased PPEs and triggered Ca^2+^ transients after steady-rate pacing with ISO incubation (*P* < 0.019) (Figure 4C). These results demonstrate that RYR2-S2814 phosphorylation in the same monomer as the CPVT pathogenic variant is required for aberrant RYR2 Ca^2+^ release at the level of individual cardiomyocytes, while RYR2-S2814 phosphorylation on wild type monomers is dispensable.

**Figure 4.**
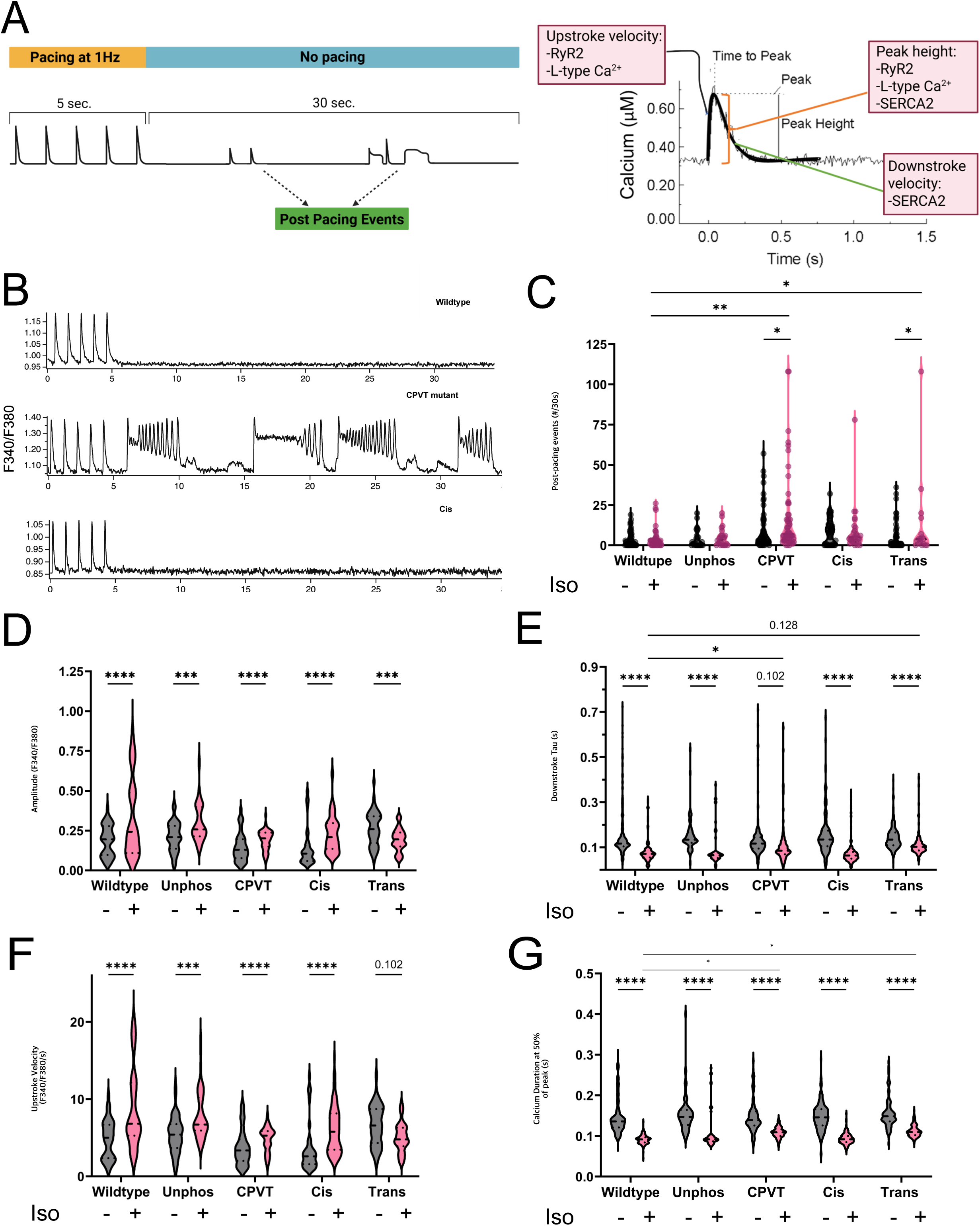
Intracellular calcium handling in RYR2 mutant cardiomyocytes under adrenergic stimulation. **A,** Cells were field-stimulated at 1 Hz for 5 seconds to record paced Ca^2+^ transients, followed by a 30-second no-pacing interval to detect spontaneous post-pacing Ca^2+^ release events. A representative Ca^2+^ transient trace depicts upstroke velocity, peak amplitude, and downstroke velocity. This panel was created using BioRender. **B,** Representative Ca^2+^ transient traces from cardiomyocytes isolated from wildtype, *CPVT mutant*, and *Cis* mice under isoproterenol (ISO) stimulation. To detect differences across genotypes and treatment conditions for Ca^2+^ transient data, we used a generalized linear model for count data (negative binomial distribution with log link) to account for non-normal, heteroscedastic data. Additional details are outlined in the methods section. **C,** Quantification of post-pacing Ca^2+^ release events (PPEs) observed during the 30-second post-pacing window, at baseline and after ISO stimulation. Significant increases in PPEs were detected in the *CPVT mutant* and *Trans* groups after ISO stimulation. There was a significant main effect of genotype (χ²(4) = 19.70, *P* < 0.001) and treatment (χ²(1) = 11.09, *P* < 0.001), though the genotype × treatment interaction did not reach statistical significance (χ²(4) = 7.37, *P* = 0.118). Post hoc Holm-adjusted contrasts revealed a significant increase in PPE from baseline to ISO in both the CPVT group (*P* = 0.032) and the Trans group (*P* = 0.026). Under ISO stimulation, both CPVT (*P* = 0.002) and Trans (*P* = 0.012) animals exhibited significantly higher PPEs compared to wildtype controls. **D,** Peak Ca^2+^ transient amplitudes at baseline and after ISO stimulation were determined by the calculating the ratio of Fura2 fluorescence (F340/F380). There was no significant main effect of genotype (χ²(4) = 6.70, *P* = 0.153), but there was a significant main effect of treatment (χ²(1) = 414.56, *P* < 0.001) and a significant genotype × treatment interaction (χ²(4) = 80.16, *P* < 0.001). There was a strong effect of ISO stimulation, with all genotypes showing significant increases in peak amplitude from baseline (Holm-adjusted *P* < 0.01). There were no statistically significant differences among genotypes at either baseline or ISO conditions after multiple-comparison correction. **E,** Single exponentials were fit to the downstroke portion of each Ca^2+^ transient, and the time constant was determined (Downstroke Tau). There was no significant main effect of genotype (χ²(4) = 3.31, P = 0.508), but a robust main effect of ISO treatment (χ²(1) = 394.19, P < 0.001) and a significant genotype × treatment interaction (χ²(4) = 63.13, P < 0.001). Holm-adjusted post hoc contrasts showed that all genotypes exhibited significantly shorter Tau values under ISO compared to baseline except for CPVT animals (P = 0.102). Conversely, CPVT animals demonstrated impaired Ca^2+^ recovery under ISO compared to wildtype (P = 0.017). **F,** Ca^2+^ transient upstroke velocity (F340/F380/s) at baseline and after ISO stimulation (100 nM). There was no significant main effect of genotype (χ²(4) = 7.15, *P* = 0.128), but there was a significant main effect of ISO treatment (χ²(1) = 383.89, *P* < 0.001) and a significant genotype × treatment interaction (χ²(4) = 81.10, *P* < 0.001). Across groups, ISO significantly increased upstroke velocity within each genotype compared to baseline, while Trans animals demonstrated a trend towards a decrease (P=0.102). **G,** Ca^2+^ transient full-width at half-maximum (CaTD_50_) was measured at baseline and with ISO stimulation. There was no significant main effect of genotype (χ²(4) = 6.66, P = 0.155), but a robust main effect of ISO treatment (χ²(1) = 1452.36, P < 0.001) and a significant genotype × treatment interaction (χ²(4) = 82.97, P < 0.001). Holm-adjusted post hoc contrasts showed that all genotypes exhibited significantly shorter CaTD_50_ values with ISO compared to baseline (all P < 0.001). Under ISO, CPVT (P = 0.041) and Trans animals (P = 0.034) had significantly prolonged CaTD_50_ values compared to wildtype controls. For all analyzed experiments the number of cells analyzed (N) were: WT: baseline = 53, ISO = 43; *Ryr2-Unphos*: baseline = 21, ISO = 13; *Ryr2-CPVT*: baseline = 34, ISO = 36; *Ryr2-Cis*: baseline = 35, ISO = 40; *Ryr2-Trans*: baseline = 51, ISO = 20. Analyses were performed using GraphPad Prism version 10.5.0. Significance values: * P<0.05, ** P<0.01, *** P <0.005, **** P <0.001.

Adrenergic stimulation induces stereotypical changes at the cardiomyocyte level to improve cardiac function and performance. To determine if the ablation of RYR2 phosphorylation by CaMKII affected adrenergic augmentation of cardiomyocyte function, we investigated Ca^2+^ transient parameters associated with β-adrenergic stimulation^13^. With pacing at 1 Hz, ISO clearly augmented peak amplitude in cardiomyocytes derived from each genotype tested (Figure 4D). While overall, adrenergic stimulation shortened the Ca^2+^ transient recovery time from peak to baseline, there was no significant shortening in the *Ryr2-CPVT* cardiomyocytes (P=0.102) (Figure 4E). ISO enhanced upstroke velocity in all but *Ryr2-Trans* cardiomyocytes (Figure 4F). Unlike the other genotypes, the baseline upstroke velocity of *Ryr2-Trans* cardiomyocytes was significantly elevated at baseline and indeed did not demonstrate any further augmentation with ISO (Figure 4F). Finally, was also a shortening of the total Ca^2+^ transient duration (CaTD_50_) as determined by the full-width at half-maximal of the recorded transients during pacing in all genotypes with ISO stimulation (Figure 4G). However, there was less shortening in response to ISO stimulation in both *Ryr2-CPVT* and *Ryr2-Trans* genotypes as compared to WT cardiomyocytes, consistent with persistently elevated diastolic Ca^2+^ levels.

Collectively, these results demonstrate that RYR2-S2814 phosphorylation in the same monomer as the RYR2-R4650I CPVT mutation is required for RYR2-R4650I to exhibit abnormal Ca^2+^ release events, while RYR2-S2814 phosphorylation (in cis, trans, or both) is dispensable for the stereotypic effects of adrenergic signaling on the cardiomyocyte Ca^2+^ transient.

### Effects of intramolecular arrangement of RYR2-S2814 phosphorylation on calcium release

To investigate the direct effects of RYR2-S2814 phosphorylation on Ca^2+^ release, we performed line-scan confocal imaging of isolated cardiomyocytes loaded with Fluo-4^22^. Without pacing, we observed spontaneous Ca^2+^ sparks with a random spatial distribution at baseline and after ISO stimulation (Figure 5A). ISO significantly increased the Ca^2+^ spark mass, and this increase was comparable across the tested genotypes (Figure 5B). In contrast, spark frequency was significantly modulated by genotype and adrenergic stimulation (Figure 5C). Baseline spark frequency was significantly higher in *Ryr2-CPVT* (*P* < 0.001) and *Ryr2-Trans* (*P* < 0.001) compared to wildtype, while *Ryr2-Cis* did not significantly differ from wild type. ISO stimulation increased spark frequency above baseline in *Ryr2-CPVT* (*P* <0.001) and *Ryr2-Trans* (*P* < 0.001), but not *Ryr2-Cis*. These data indicate the diastolic RYR2 opening caused by the CPVT variant requires RYR2-S2814 phosphorylation on the same monomer. That *Ryr2-CPVT* and *Ryr2-Trans* but not *Ryr2-Cis* exhibited elevated frequency of baseline channel opening suggests some degree of RYR2-S2814 phosphorylation occurs without ISO stimulation.

**Figure 5.**
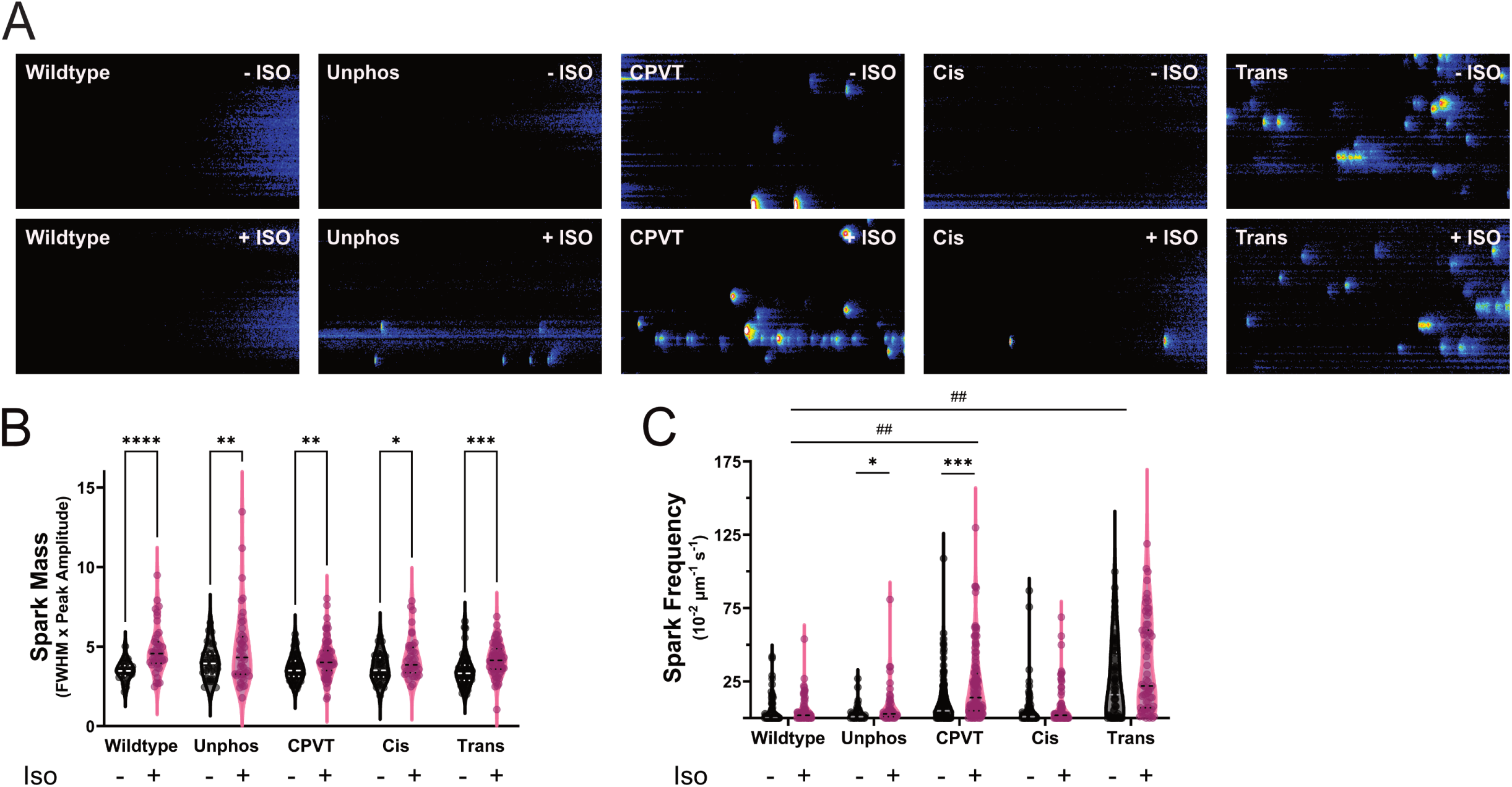
Calcium spark activity in RYR2 mutant cardiomyocytes at baseline and under adrenergic stimulation. **A,** Representative line-scan confocal images of Ca^2+^ sparks in cardiomyocytes isolated from wildtype, *Unphos*, *CPVT mutant*, *Cis*, and *Trans* mice, shown at baseline and after isoproterenol stimulation (100 nM). The traces illustrate differences in spontaneous Ca^2+^ release activity across genotypes and conditions. **B,** Isoproterenol stimulation significantly increased spark mass in all groups, consistent with enhanced RYR2-mediated Ca^2+^ release under adrenergic conditions. Two-way ANOVA: treatment (baseline vs. iso): F(1, 495) = 49.13, *P* < 0.001, genotype: F(4, 495) = 3.207, *P* = 0.0129, genotype-by-treatment: F(4, 495) = 1.140, *P* = 0.3367. Post hoc test: Tukey. **C,** Spark frequency measured at baseline and after isoproterenol stimulation. * Indicates significance of spark frequency before and after isoproterenol stimulation. # indicates significance between both groups of wild-type cells compared to other tested genotypes. Spark frequency was significantly elevated in the *CPVT* mutant and *Trans* groups following isoproterenol. A generalized linear model for count data (negative binomial distribution with log link) revealed a significant main effect of genotype (χ²(4) = 115.96, *P* < 0.001) and treatment (χ²(1) = 17.27, *P* < 0.001), while the genotype × treatment interaction showed a statistical trend (χ²(4) = 5.24, *P* = 0.263). Post hoc contrasts (Holm-adjusted, α = 0.05) revealed significant increase in spark frequency from baseline to ISO in the Unphos group (IRR = 2.46, 95% CI [1.25–4.85], *P* = 0.038) and the CPVT group (IRR = 2.07, 95% CI [1.39–3.07], *P* = 0.002). At Baseline, both CPVT and Trans animals displayed elevated spark frequency relative to WT (CPVT IRR = 0.48, 95% CI [0.30–0.78], *P* = 0.0028; Trans IRR = 0.22, 95% CI [0.13–0.37], *P* = 9.4×10⁻⁹). Under Iso stimulation, spark frequency was significantly higher in CPVT compared to WT (IRR = 0.27, 95% CI [0.17–0.44], *P* < 0.001) and in Trans compared to WT (IRR = 0.17, 95% CI [0.10–0.29], *P* < 0.001). All analyses were performed using GraphPad Prism version 10.5.0. Significance values: * P<0.05, ** P<0.01, *** P <0.005, **** P <0.001.

## Discussion

Inherited arrhythmias such CPVT, caused predominantly by variants in RYR2 that increase channel open probability, show how discrete molecular lesions can provoke catastrophic electrical instability in structurally normal hearts during adrenergic stress^23–25^. Despite two decades of investigation, the field has lacked clarity on how the spatial arrangement of mutations and regulatory phosphorylation sites within the RYR2 tetramer contributes to channel dysfunction. Our study addresses this knowledge gap using allele-specific mouse models that precisely manipulate the structural configuration of a disease-causing mutation (R4650I) and a regulatory phosphorylation-null variant (S2814A), offering in vivo evidence that RYR2 function is modulated at the monomer level.

By comparing *Cis* and *Trans* configurations—placing S2814A and R4650I on the same or opposing alleles, respectively—we demonstrate that functional rescue requires that the CPVT-causing mutation and S2814 regulatory ablation occur within the same monomer. Only the *Cis* model, in which the S2814A mutation resides on the same monomer as R4650I, exhibited complete suppression of arrhythmogenic Ca^2+^ release, spontaneous post-pacing activity, and ventricular tachycardia. In contrast, the *Trans* model, despite harboring the same mutations, failed to suppress Ca^2+^ sparks (Figure 5), post-pacing events (Figure 4), or arrhythmia at the whole organ level (Figure 3), emphasizing the necessity of the regulatory ablation within the same monomer as the CPVT mutation. These findings suggest that phosphorylation at S2814 acts locally to modulate the function of the specific RYR2 monomer in which it resides, and that therapeutic benefit is only achieved when phosphorylation site ablation and pathogenic mutation are conformationally aligned within the same protomer.

While bradycardia is a clinical feature in up to 50% of CPVT patients, the exact mechanism is incompletely understood^7^. Unexpectedly, the regulatory-null mutant (RYR2-Unphos) or the cis-configuration of S2814A and R4650I did not produce the same heart regulation defects seen with either the *RYR2-CPVT* or *RYR2-Trans* models. These data suggest that the impairment in heart rate regulation is precipitated by the presence of a GOF RYR2 variant but is effectively mitigated by the inhibition of phosphorylation at S2814 on the same monomer. This further supports the hypothesis that any molecular changes induced by S2814 phosphorylation that unmasks the effects of GOF RYR2 variants channel activity is universal and is present in both the working ventricular myocardium and Sino-atrial node. In contrast, we did not observe any significant defects in Ca^2+^ regulation secondary to adrenergic stimulation at the single-cell level in our different animal models expect for the lack of change in upstroke velocity in the *RYR2-Trans* cardiomyocytes (Figure 4). This suggests that the Trans configuration may represent a partially stimulated state consistent with the blunted heart rate responses to isoproterenol and epinephrine (Figure 3). Ultimately these data suggest that the inhibition of S2814 phosphorylation can reverse the physiologic effects of CPVT RYR2 variants but does not impair normal physiologic effects of β-adrenergic stimulation provided an attractive target for selective anti-arrhythmic therapy.

This monomer-specific effect helps resolve a long-standing and contentious debate surrounding RYR2 phosphorylation. In the early 2000s, the CPVT-RYR2 field focused heavily on a highly influential model implicating PKA-mediated phosphorylation at S2808 as a central mechanism of RYR2 dysfunction in heart failure and arrhythmia^18,26^. According to this model, chronic sympathetic stimulation leads to hyperphosphorylation of S2808, dissociation of the stabilizing protein FKBP12.6, and aberrant RYR2 opening. While foundational, this theory has since been challenged by several groups who failed to reproduce the findings, observed minimal phenotypes in S2808A knock-in models, or demonstrated compensatory mechanisms involving other phosphorylation sites^27,28^. In contrast, a growing body of literature has pointed to CaMKII-dependent phosphorylation at S2814 as a more potent and arrhythmogenic modulator of RYR2 activity^10,29,30^.

Our findings provide strong support for this shift in focus. In our allele-specific models, S2808 and S2030 remained phosphorylatable and functionally intact across genotypes, confirming that global adrenergic signaling was unchanged. Moreover, the inability of S2808 phosphorylation to compensate in the *Trans* model, combined with the dominant rescue effect seen in the *Cis* configuration, strongly position S2814 as the critical pathogenic node, which has significant implications for both basic mechanisms and targeted therapeutic design. Also intriguing is that in both genotypes without any arrhythmia including *Ryr2-Unphos* and *Ryr2-Cis* there was actually increased phosphorylation at S2030 which has been associated with a protective effect for RYR2-dependent arrhythmia and Ca^2+^ alternans^31^. Yet, multiple other regulatory mechanisms influence RYR2 gating, including luminal Ca^2+^ sensors (e.g., calsequestrin, triadin) and redox modifications (e.g., oxidation, S-nitrosylation)^21,32^, but our findings indicate that these processes are not sufficient to override the requirement for CaMKII-mediated phosphorylation at S2814 in CPVT pathogenesis. These results support the conclusion that phosphorylation at S2814 is both necessary and non-redundant for the pathological phenotype in our model, highlighting a critical intra-monomeric mechanism of RYR2 dysregulation.

A unique strength of this study lies in its systematic investigation of RYR2 function across multiple biological scales: whole-animal electrophysiology, isolated cardiomyocyte calcium imaging, and biochemical analysis of individual channel modifications. The remarkable consistency of the results across these three domains significantly enhances the robustness and translational relevance of our findings.

### Limitations

The primary limitation of this study is the inability to resolve the stoichiometry of RYR2 tetramers in the *Trans* model. Although the two mutations were engineered on opposite alleles, monomer assembly is stochastic, and individual channels may consist of variable combinations of mutant and non-mutant monomers. This channel-level heterogeneity may partially explain the intermediate phenotypes observed in some assays. Nonetheless, the persistent failure of the *Trans* model to suppress arrhythmia, despite harboring protective monomers, strongly supports the conclusion that functional rescue requires phosphorylation ablation within the same monomer as the disease-causing mutation. Future studies using super-resolution imaging techniques, such as stochastic optical reconstruction microscopy (STORM) or stimulated emission depletion (STED), could provide the spatial resolution necessary to directly visualize monomer arrangement and composition. Additionally, advanced mass spectrometry approaches, including cross-linking mass spectrometry (XL-MS), may help identify and quantify specific phosphorylation events and inter-monomer interactions at the molecular level, thereby overcoming current experimental limitations. Another limitation is that we only studied the incorporation of the S2814A mutation with one CPVT variant.

## Conclusions

This study provides the first in vivo evidence that spatial co-localization of pathogenic RYR2 mutations and regulatory phosphorylation sites within individual monomers is a critical determinant of arrhythmia susceptibility. By leveraging genetic tools to define monomer-specific configurations, we uncover a previously unappreciated mechanism of RYR2 regulation and clarify conflicting findings in the literature. These findings open a new therapeutic dimension for CPVT and related disorders, where allele-specific or monomer-targeted interventions may offer greater efficacy than global modulation. Ultimately, this work highlights the importance of structure-informed molecular design in the treatment of inherited arrhythmia syndromes.

## Notes

### Competing Interest Statement

The authors have declared no competing interest.

## References

1. Gorski PA, Ceholski DK, Hajjar RJ. Altered myocardial calcium cycling and energetics in heart failure--a rational approach for disease treatment. Cell Metab. 2015;21:183–194.

2. Marks AR. Targeting ryanodine receptors to treat human diseases. J Clin Invest [Internet]. 2023;133. Available from: https://www.jci.org/articles/view/162891

3. Woll KA, Van Petegem F. Calcium-release channels: structure and function of IP3 receptors and ryanodine receptors. Physiol Rev. 2022;102:209–268.

4. Dobrev D, Wehrens XHT. Role of RyR2 phosphorylation in heart failure and arrhythmias: Controversies around ryanodine receptor phosphorylation in cardiac disease. Circ. Res. 2014;114:1311–9; discussion 1319.

5. Lanner JT, Georgiou DK, Joshi AD, Hamilton SL. Ryanodine receptors: structure, expression, molecular details, and function in calcium release. Cold Spring Harb Perspect Biol. 2010;2:a003996.

6. Venetucci L, Denegri M, Napolitano C, Priori SG. Inherited calcium channelopathies in the pathophysiology of arrhythmias. Nat Rev Cardiol. 2012;9:561–575.

7. Miyata K, Ohno S, Itoh H, Horie M. Bradycardia Is a Specific Phenotype of Catecholaminergic Polymorphic Ventricular Tachycardia Induced by RYR2 Mutations. Intern Med. 2018;57:1813– 1817.

8. Park S-J, Zhang D, Qi Y, Li Y, Lee KY, Bezzerides VJ, Yang P, Xia S, Kim SL, Liu X, Lu F, Pasqualini FS, Campbell PH, Geva J, Roberts AE, Kleber AG, Abrams DJ, Pu WT, Parker KK. Insights Into the Pathogenesis of Catecholaminergic Polymorphic Ventricular Tachycardia From Engineered Human Heart Tissue. Circulation. 2019;140:390–404.

9. Wehrens XHT, Lehnart SE, Reiken S, Vest JA, Wronska A, Marks AR. Ryanodine receptor/calcium release channel PKA phosphorylation: a critical mediator of heart failure progression. Proc Natl Acad Sci U S A. 2006;103:511–518.

10. Wehrens XHT, Lehnart SE, Reiken SR, Marks AR. Ca2+/calmodulin-dependent protein kinase II phosphorylation regulates the cardiac ryanodine receptor. Circ Res. 2004;94:e61–70.

11. Bezzerides VJ, Caballero A, Wang S, Ai Y, Hylind RJ, Lu F, Heims-Waldron DA, Chambers KD, Zhang D, Abrams DJ, Pu WT. Gene Therapy for Catecholaminergic Polymorphic Ventricular Tachycardia by Inhibition of Ca2+/Calmodulin-Dependent Kinase II. Circulation. 2019;140:405– 419.

12. Gao S, Ho D, Vatner DE, Vatner SF. Echocardiography in Mice. Curr Protoc Mouse Biol. 2011;1:71–83.

13. O’Connell TD, Rodrigo MC, Simpson PC. Isolation and culture of adult mouse cardiac myocytes. Methods Mol Biol. 2007;357:271–296.

14. Cheng H, Song LS, Shirokova N, González A, Lakatta EG, Ríos E, Stern MD. Amplitude distribution of calcium sparks in confocal images: theory and studies with an automatic detection method. Biophys J. 1999;76:606–617.

15. Schneider L, Begovic M, Zhou X, Hamdani N, Akin I, El-Battrawy I. Catecholaminergic polymorphic ventricular tachycardia: Advancing from molecular insights to preclinical models. J Am Heart Assoc. 2025;14:e038308.

16. Napolitano C, Bloise R, Memmi M, Priori SG. Clinical utility gene card for: Catecholaminergic polymorphic ventricular tachycardia (CPVT). Eur J Hum Genet [Internet]. 2014;22. Available from: 10.1038/ejhg.2013.55

17. Leenhardt Antoine, Denjoy Isabelle, Guicheney Pascale. Catecholaminergic Polymorphic Ventricular Tachycardia. Circ Arrhythm Electrophysiol. 2012;5:1044–1052.

18. Marx SO, Reiken S, Hisamatsu Y, Jayaraman T, Burkhoff D, Rosemblit N, Marks AR. PKA Phosphorylation Dissociates FKBP12.6 from the Calcium Release Channel (Ryanodine Receptor). Cell. 2000;101:365–376.

19. Shinoda Y, Komatsu Y, Nogami A, Igarashi M, Yamasaki H, Sekiguchi Y, Aonuma K, Ieda M. Stepwise approach to induce infrequent premature ventricular complex using bolus isoproterenol and epinephrine infusion. Pacing Clin Electrophysiol. 2020;43:437–443.

20. Roston TM, Vinocur JM, Maginot KR, Mohammed S, Salerno JC, Etheridge SP, Cohen M, Hamilton RM, Pflaumer A, Kanter RJ, Potts JE, LaPage MJ, Collins KK, Gebauer RA, Temple JD, Batra AS, Erickson C, Miszczak-Knecht M, Kubuš P, Bar-Cohen Y, Kantoch M, Thomas VC, Hessling G, Anderson C, Young M-L, Cabrera Ortega M, Lau YR, Johnsrude CL, Fournier A, Kannankeril PJ, Sanatani S. Catecholaminergic polymorphic ventricular tachycardia in children: analysis of therapeutic strategies and outcomes from an international multicenter registry. Circ Arrhythm Electrophysiol. 2015;8:633–642.

21. Wleklinski MJ, Kannankeril PJ, Knollmann BC. Molecular and tissue mechanisms of catecholaminergic polymorphic ventricular tachycardia. J Physiol. 2020;598:2817–2834.

22. Bray M-A, Geisse NA, Parker KK. Multidimensional detection and analysis of Ca2+ sparks in cardiac myocytes. Biophys J. 2007;92:4433–4443.

23. Priori SG, Napolitano C, Memmi M, Colombi B, Drago F, Gasparini M, DeSimone L, Coltorti F, Bloise R, Keegan R, Cruz Filho FES, Vignati G, Benatar A, DeLogu A. Clinical and molecular characterization of patients with catecholaminergic polymorphic ventricular tachycardia. Circulation. 2002;106:69–74.

24. Cerrone M, Napolitano C, Priori SG. Catecholaminergic polymorphic ventricular tachycardia: A paradigm to understand mechanisms of arrhythmias associated to impaired Ca(2+) regulation. Heart Rhythm. 2009;6:1652–1659.

25. Laitinen PJ, Brown KM, Piippo K, Swan H, Devaney JM, Brahmbhatt B, Donarum EA, Marino M, Tiso N, Viitasalo M, Toivonen L, Stephan DA, Kontula K. Mutations of the cardiac ryanodine receptor (RyR2) gene in familial polymorphic ventricular tachycardia. Circulation. 2001;103:485–490.

26. Wehrens XHT, Lehnart SE, Reiken S, van der Nagel R, Morales R, Sun J, Cheng Z, Deng S-X, de Windt LJ, Landry DW, Marks AR. Enhancing calstabin binding to ryanodine receptors improves cardiac and skeletal muscle function in heart failure. Proc Natl Acad Sci U S A. 2005;102:9607– 9612.

27. Houser SR. Role of RyR2 phosphorylation in heart failure and arrhythmias: protein kinase A-mediated hyperphosphorylation of the ryanodine receptor at serine 2808 does not alter cardiac contractility or cause heart failure and arrhythmias. Circ Res. 2014;114:1320–7; discussion 1327.

28. Benkusky NA, Farrell EF, Valdivia HH. Ryanodine receptor channelopathies. Biochem Biophys Res Commun. 2004;322:1280–1285.

29. Respress JL, van Oort RJ, Li N, Rolim N, Dixit SS, deAlmeida A, Voigt N, Lawrence WS, Skapura DG, Skårdal K, Wisløff U, Wieland T, Ai X, Pogwizd SM, Dobrev D, Wehrens XHT. Role of RyR2 phosphorylation at S2814 during heart failure progression. Circ Res. 2012;110:1474–1483.

30. Terentyev D, Nori A, Santoro M, Viatchenko-Karpinski S, Kubalova Z, Gyorke I, Terentyeva R, Vedamoorthyrao S, Blom NA, Valle G, Napolitano C, Williams SC, Volpe P, Priori SG, Gyorke S. Abnormal interactions of calsequestrin with the ryanodine receptor calcium release channel complex linked to exercise-induced sudden cardiac death. Circ Res. 2006;98:1151–1158.

31. Wei J, Guo W, Wang R, Paul Estillore J, Belke D, Chen Y-X, Vallmitjana A, Benitez R, Hove-Madsen L, Chen SRW. RyR2 Serine-2030 PKA site governs Ca ^2+^ release termination and Ca ^2+^ alternans. Circ Res [Internet]. 2023 [cited 2025 Jul 12];132. Available from: 10.1161/CIRCRESAHA.122.321177

32. Terentyev D, Györke I, Belevych AE, Terentyeva R, Sridhar A, Nishijima Y, de Blanco EC, Khanna S, Sen CK, Cardounel AJ, Carnes CA, Györke S. Redox modification of ryanodine receptors contributes to sarcoplasmic reticulum Ca2+ leak in chronic heart failure. Circ Res. 2008;103:1466–1472.

